# An intergeneric recombinant geminivirus causes soybean stay-green disease

**DOI:** 10.1101/2022.02.09.479675

**Authors:** Ruixiang Cheng, Ruoxin Mei, Rong Yan, Hongyu Chen, Dan Miao, Lina Cai, Jiayi Fan, Gairu Li, Ran Xu, Weiguo Lu, Yu Gao, Wenwu Ye, Shuo Su, Tianfu Han, Junyi Gai, Yuanchao Wang, Xiaorong Tao, Yi Xu

**Affiliations:** Department of Plant Pathology, Nanjing Agricultural University, Nanjing 210095, China; The Key Laboratory of Biological Interaction and Crop Health, Nanjing Agricultural University, Nanjing 210095, China; Engineering Laboratory of Animal Immunity of Jiangsu Province, College of Veterinary Medicine, Academy for Advanced Interdisciplinary Studies, Nanjing Agricultural University, Nanjing 210095, Jiangsu, China; Crop Research Institute, Shandong Academy of Agricultural Sciences, Jinan 250100, Shandong, China; Henan Academy of Crop Molecular Breeding, Zhengzhou 450002, Henan, China; College of Plant Protection, Jilin Agricultural University, Changchun 130018, Jilin, China; MOA Key Laboratory of Soybean Biology (Beijing), Institute of Crop Sciences, Chinese Academy of Agricultural Sciences, Beijing 100081, China; Soybean Research Institute, Nanjing Agricultural University, Nanjing 210095, Jiangsu, China

**Author notes:** **Correspondence:** X.T, Y.X.

## Abstract

Soybean is one of the most valuable legume crops in the world with high nutritional value. Recently, the outbreak of soybean stay-green syndrome has swept the soybean production in the Huang-Huai-Hai region of China, resulting in huge yield losses, which has become an epidemic and prominent problem in soybean production. However, the cause of the stay-green syndrome remains obscure. Here, we report a novel intergeneric recombinant geminivirus which causes soybean stay-green symptoms. Viral small RNA-based screening identified a new recombinant geminvirus from field soybean stay-green samples. The complete genome sequence of the virus contains 2762 nucleotide (nt) and appears to be an intergeneric recombinant virus in which protein coding for coat protein (V1) is similar to member of genus *Mastrevirus*, whereas proteins coding for V2, C2, C3 are most similar to those of viruses in the *Maldovirus* genus, and C1 and C4 are most similar to virus in genus *Begomovirus*. Inoculation of the infectious clone of the recombinant geminivirus through *Agrobacterium rhizogenes* causes typical soybean stay-green syndrome which resembles field symptoms including delayed leaf senescence, flat pods and abnormal seeds. The recombinant geminivirus can be detected in seed coat but not in cotyledon and embryo, thus failing to be transmitted by seeds. Moreover, the genome variation and epidemiological dynamic analysis were also carried out to help the continuous epidemiological surveillance of this emerging geminivirus. Collectively, this new geminivirus is tentatively named soybean stay-green associated virus (SoSGV). Our determination of the causal agent of soybean stay-green disease will bolster efforts to develop effective management strategies to control this prevalent disease in the field.

---

Plant viruses make up almost half of the plant disease-causing pathogens, affecting crop yields and the global economy (Savary et al., 2019). Soybean [*Glycine max* (L.) Merr.] is one of the most valuable legume crops in the world, supplying 25% of the global edible oil and two-thirds of the global concentrated protein for livestock feeding. Recently, the outbreak of soybean stay-green syndrome with delayed leaf senescence (stay-green), flat pods and increased number of abnormal seeds has swept the soybean production in the Huang-Huai-Hai region of China, resulting in huge yield losses (Xu et al., 2019). This disease has become an epidemic and prominent problem in soybean production and is still expanding its geography, including North America, posing a serious threat to soybean production (Harbach et al., 2016; Li et al., 2019; Zhang et al., 2016). However, the cause of the stay-green syndrome remains obscure.

To investigate the possible pathogenic agents associated with this stay-green syndrome, soybean samples with typical symptoms were collected from several independent fields in Huang-Huai-Hai region of China (*SI Appendix* Fig. S1 *A-C*). Those samples were mixed and identified with small RNA-based screening. Of the 66,212 contigs assembled by Velvet (v1.2.10), one contig with a length of 311 nucleotides (nt) showed the high sequence identity (89.1%) to a begomovirus C1 (tomato leaf curl Java virus-[Ageratum], TLCJV, Genbank No. AB162141.1). The complete viral genomic sequence was further determined to be 2,762 nucleotides (nt) in length. There were two ORFs from virion strand and four ORFs from complementary-sense (Fig. 1*A*). The size of the intergenic region (IR) was 283 nt which is similar to that of most begomoviruses (200-300 nt) (Hanley-Bowdoin et al., 2013). A nonanucleotide sequence (‘TAATATTAC’) was found in the viral intergenic region (*SI Appendix*, Fig. S1*D*) and the IR of this geminivirus contains TATA-boxes in the virion (position 93) and the complementary sense transcripts (position 2694). An iterated DNA motif (iteron) (GGAGA, positions 2673-2677) and the corresponding iteron-related domain (IRD) showing the amino acid motif of PRRFRIQ in the N-terminal of C1 (Rep) were identified (*SI Appendix* Fig. S1*E*), indicating this virus employs a typical begomovirus-specific replication strategy (Arguello-Astorga and Ruiz-Medrano, 2001). This virus was tentatively designated as soybean stay-green associated virus (SoSGV) based on the symptoms of soybean found in the field. The whole genome sequence of SoSGV has been assigned GenBank accession number OM145986. Moreover, PCR assays were performed using a pair of primer designed to amplify the full-length CP (V1), leading to the detection of SoSGV in 67.5% of 40 samples collected in two years from seven fields across two provinces in the Huang-Huai-Hai region of China (Fig. 1*B*). All 27 samples with positive SoSGV showed stay-green symptoms including delayed leaf senescence, flat pods and abnormal seeds (Fig. 1*B*), which indicated that SoSGV was highly correlated with soybean stay-green syndrome in the Huang-Huai-Hai region of China.

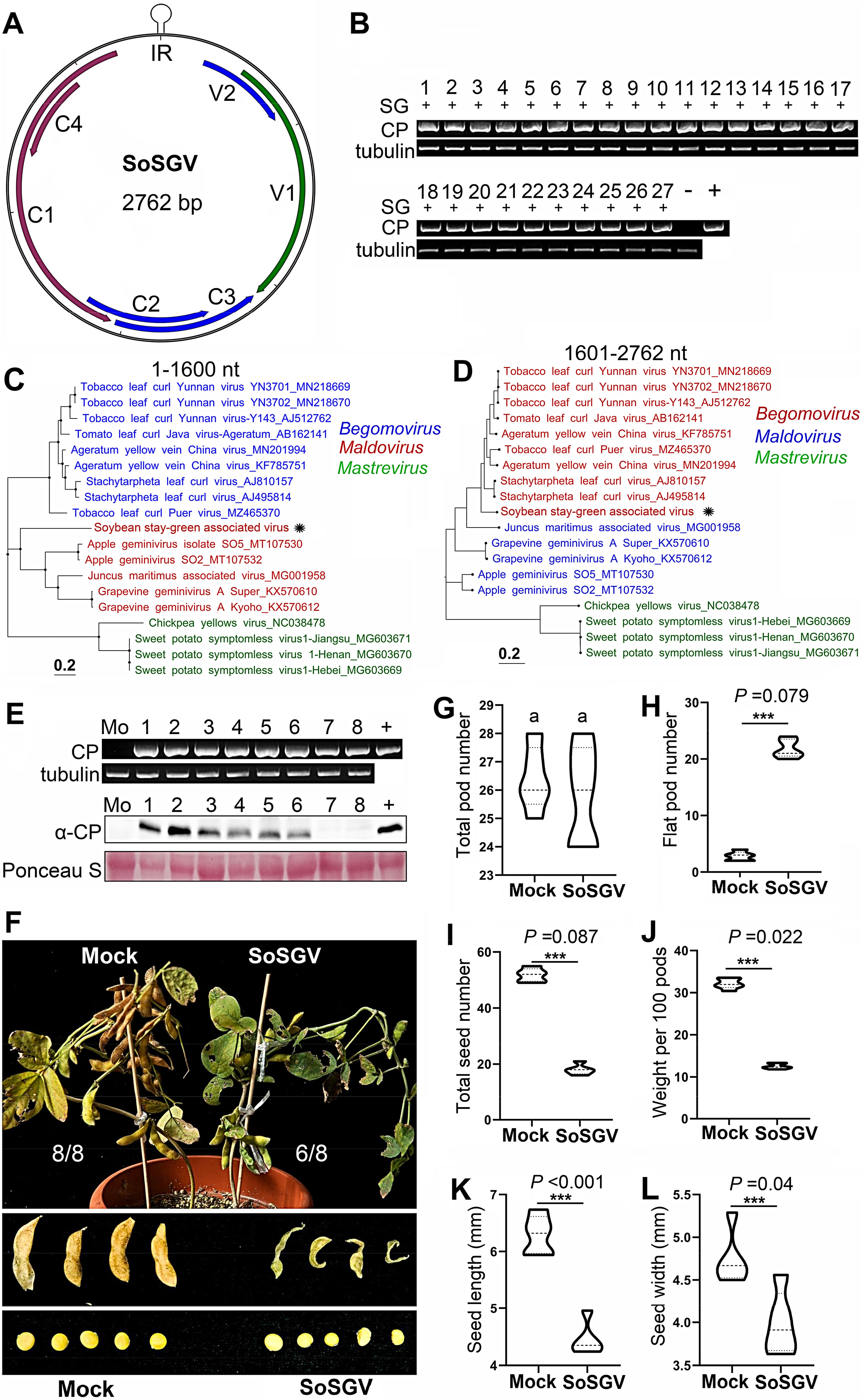
Determination of SoSGV is the causal agent of soybean stay-green disease. (*A*), The genome structure of soybean stay-green associated virus (SoSGV). (*B*), Detection of SoSGV in 27 samples collected in two years from seven fields across two provinces in the Huang-Huai-Hai region of China, and all these samples showing stay-green symptom. SG, stay-green symptom including delayed leaf leaf senescence, empty pods and abnormal seeds. “−”, field sample without stay-green symptom; “+”, a plasmid containing pBin2x-SoSGV used as a positive control, and soybean *tubulin* was used as the internal gene. The numbers 1-27 represent samples collected from four locations in Huang-Huai-Hai region of China as indicated in *SI Appendix* Table S5. Maximum-likelihood phylogenetic trees inferred for the 1-1600 nt (*C*) and 1601-2762 nt (*D*) regions. SoSGV is indicated by red text and asterisk; The branch lengths reflect the number of nucleotide substitutions per site. (*E*) The infection of SoSGV in eight affected soybean plants were confirmed by both PCR and Western blot. Plasmid pBin2x-SoSGV and the protein extract from SoSGV infected *N. benthamiana* plant were used as positive controls in PCR and Western blot, respectively. Soybean *tubulin* gene was used as an internal reference gene. (*F*) Four months after inoculation of soybean plants by infectious clone of SoSGV, all mock inoculated plants (8/8) display yellowing leaves with mature seeds, whereas for SoSGV infection, the leaves of six out eight plants are still stay-green and the seed development are severely affected. (*G-L*), The effect of pods number (*G*), flat pods (*H*), seeds number (*I*), weight per 100 pods (*J*), seed length (*K*) and width (*L*) after SoSGV infection. Statistical significance is indicated by different letters (*P* < 0.01). Error bars represent means ± standard deviation. Statistical analysis was performed using one sample Student t-test. *P* values<0.05 were considered significant.

Analysis of viral small interference (vsi) RNA libraries showed that most vsiRNAs were 18 to 24-nt in length and among them the 21-nt class was the most dominant class, accounting for 57.6% of vsiRNAs in the sample (*SI Appendix* Fig. S1*F*). The hotspots of the vsiRNAs appeared to be derived unevenly from SoSGV genome, with the most peaks were from virion position nt 1236, and complementary-strand position at 1298, and the peak position was near the end of C2 ORF (*SI Appendix* Fig. S1*G*).

The full-length genome sequence of SoSGV was blast against nt/nr database in NCBI. We found that nt 1601-2762 and the following 22 nt in the intergenic region showed the highest sequence identities (84%) with TLCJV (AB162141.1) which is a typical member of geminiviruses in the genus *Begomovirus*, and this region contains C1 (Rep), C4 ORFs and IR (*SI Appendix*, Table S1). Intriguingly, the other region around 1580 nt containing four ORFs including V2, V1, and C2 and C3 showed no nt similarity with any sequences in the database. Evolutionary relationship between SoSGV genome sequences and the other reference geminiviruses was analyzed and the results showed fourteen major clades. SoSGV clustered separately from any genus in the family *Geminiviridae* and it clustered mostly close to the maldoviruses in the *Maldovirus* genus but in an unclassified group (*SI Appendix*, Fig. S2). Next, similarity plots and segmented sequence phylogenetic analyses revealed that SoSGV was recombinant, including genomic fragments from maldovirus lineages (nt 23-1600) and begomovirus lineages (nt 1601-2672, and the following nt 1-22 in the IR region) (Fig. 1 *C-D*; *SI Appendix*, Fig. S3*A*). In supporting of this, ML phylogenetic analyses based on the deduced amino acid (AA) sequences of each ORF were performed. V1 (CP) showed that it was clustered most closely to V1 of sweet potato symptomless virus 1 (AWB97033.1, 29.6% AA identity) and chickpea yellows virus (YP_009506576.1, 27.2% AA identity) in the *Mastrevirus* genus (*SI Appendix*, Fig. S3*B* and Table S1). The ML phylogenetic analyses based on the AA sequences of V2, C2, and C3 showed they clustered with viruses in the *Maldovirus* genus (*SI Appendix*, Fig. S3*C* and *3E-F*). For C1 and C4, they clustered together in the begomovirues group (*SI Appendix*, Fig. S3*D* and *G*). All these findings indicate that SoSGV to be a highly divergent geminivirus and was formed by large-fragment recombinations of viruses from three genera of geminiviruses.

To determine whether the recombinant virus is the causal agent of soybean stay-green disease, an infectious clone pBin2x-SoSGV was constructed using binary vector pBinPLUS (Cui et al., 2004). The infectivity of pBin2x-SoSGV by *Agrobacterium* (GV3101)-mediated injection to a number of plant species is summarized in *SI Appendix*, Table S2. Collectively, SoSGV can infect *N. benthamiana*, *N. glutinosa* and *Datura stramonium* but not tomato varieties Moneymaker, M82 and soybean cultivars including Williams 82, Wanhua 518 and Shanning through *Agrobacterium* (GV3101)-mediated inoculation (*SI Appendix*, Fig. S4 and Table S2). Next, plasmid of pBin2x-SoSGV was transformed into *A. rhizogenes* strain K599 which can effectively infect soybean root (Veena and Taylor, 2007), and the plantlets of soybean were inoculated by *A. rhizogenes* K599 containing pBin2x-SoSGV. The results showed that >90% inoculation efficiency of SoSGV to soybean cultivar Williams 82, Wandou 518, Sanning and Nannong1138-2 mediated by *A. rhizogenes* K599 (*SI Appendix*, Fig. S5 and Table S2).

Four months after inoculation of SoSGV, all the leaves of eight soybean plants with mock treatment became yellowing with mature pods and seeds, whereas for SoSGV-inoculated plants, all eight plants were tested positive for SoSGV by PCR, but only 6 out of 8 plants could detect SoSGV CP accumulation by Western blot (Fig. 1*E*). These six positive plants still kept stay-green symptoms, including delayed leaf senescence (stay-green), increased numbers of abnormal seeds, many flat pods (Fig. 1*F*). Symptoms of plants infected with pBin2x-SoSGV were indistinguishable from symptoms produced by plants found in the field. Furthermore, the exact effects of SoSGV on seeds yield per plant were counted. The results showed that infections by using cloned SoSGV had little effect on pod number, but significantly increased flat pod number, reduced total seed number, decreased weight per hundred pods and seed size (Fig. 1 *G-L*). All these effects were consistent with the stay-green soybean found in the field. Moreover, we found that SoSGV could also cause stay-green like symptoms on *N. benthamina* plant after inoculation of pBin2x-SoSGV, including delayed leaf senescence and more empty pods of plants in comparison to plants inoculated with pBinPLUS control at 50 dpi (*SI Appendix*, Fig. S6). All these results support that SoSGV can cause plant stay-green symptoms.

The outbreak of SoSGV associated with the presence of high number of whitefiles *Bemisia tabaci* (Gennadius) and other insects in the field suggests it is transmitted by insect vector(s). Our repeated results showed that SoSGV (SoSGV-YY1) failed to be transmitted by *B. tabaci* (*SI Appendix*, Table S3). The seed-borne nature of SoSGV was then investigated and the presence of SoSGV in whole seed was detected (108 out of 120 seeds collected from field infected samples) (*SI Appendix*, Table S4). Moreover, the percentage of samples which are positive for the presence of SoSGV in seed coat dissected from the seed of infected plant is 100 (108 out of 108), however, SoSGV was never detected in endosperm and embryo (*SI Appendix*, Table S4). Next, seed transmission of SoSGV was further tested through grow-out test. Results showed that all seedlings from SoSGV-infected seeds were negative (93 germinated out of 100 planted) (*SI Appendix*, Table S4). To eliminate other factors in the field that may affect SoSGV entering seed, the presence of SoSGV in seed coat but not in cotyledon and embryo of plants inoculated by *A. rhizogenes* K599 containing pBin2x-SoSGV were confirmed by PCR (*SI Appendix*, Fig. S7*A* and Table S4). In addition, semi-quantitative PCR showed that the abundance of SoSGV in soybean seeds compared with other plant tissues (*SI Appendix*, Fig. S7*B*).

Next, complete genomes of twenty-seven SoSGV from infected soybean plants with stay-green symptoms collected in four different regions in Huang-Huai-Hai region of China from 2019 to 2021 were determined, ranging from 2760 to 2762 bp in length, and the sequences have been deposited in GenBank (*SI Appendix*, Table S5). They shared the highest nucleotide identities with a SoSGV genomic sequence collected in 2016 from South Korea, at 98.5-99.5% (Dataset S1). High percent identities of all encoded proteins among all isolates from different geographic origin were observed (Dataset S1). Selection pressures analysis showed that only one codon (amino acid position 219 in C1) was positively selected among all SoSGV-encoded proteins, indicating that SoSGV is mainly undergoing purified selection currently (*SI Appendix*, Table S6). Next, the molecular variability was further examined based on Bayesian and MCC analysis. The complete SoSGV genomes from soybean fields formed a monophyletic group comprising two relatively divergent sister clades (I and II), while the SoSGV genotypes from South Korea clustered as closely related clade I (*SI Appendix*, Fig. S8). MCC analysis and the ML analysis showed that the geographical distribution of SoSGV isolates was not completely correlated, which indicated that the geographical correlation of SoSGV isolates was not significant (*SI Appendix*, Fig. S8 and Dataset S1).

To better understand the origin, evolution and epidemiological dynamics of SoSGV, the time of most recent ancestor (tMRCA) and nucleotide substitution rate of the complete genomic sequences was estimated by using BEAST (v1.10.4). The tMRCA of SoSGV was 1911.75 (95% HPD:1821.8-1998.5), individually, the clade I was in 1930.57 (95% HPD:1853.12-2002.22), and the clade II was in 1946.85 (95% HPD:1870.99-2004.48) (*SI Appendix*, Fig. S8). Additionally, the mean substitution rate of the whole genome sequences of all SoSGV isolates was 1.2×10^−4^ (95% HPD: 3.11×10^−5^-2.8×10^−4^) substitution/site/year calculated by BEAST software, which was close to other reported geminiviruses (Duffy and Holmes, 2008). And the substitution rate for clade I and clade II were 1.15×10^−4^ (95% HPD: 2.33×10^−5^-2.61×10^−4^) and 1.32×10^−4^ (95% HPD:3.56×10^−5^-3.1×10^−4^) substitution/site/year, respectively (Dataset S2).

In conclusion, we determine the etiology of an emerging soybean stay-green disease in China. The causal agent is a newly evolved geminivirus formed by large-fragment intergeneric recombination from genera in the family of *Geminiviridae*. The accumulation of a large amount of SoSGV in the seed coat and the stagnation of seed development after SoSGV infection indicate SoSGV may hinder the transportation of photoassimilates from vegetative tissues including maternal seed coat to cotyledon and embryo, leading to imbalance of source-sink partitioning in favor of the source, thus delaying the senescence process. Our study supplies both biological properties and epidemiological characteristics of SoSGV and the determination of the causal agent of soybean stay-green disease will bolster efforts to develop effective management strategies to control this soybean epidemic.

## Supporting information

All supplemental materials

## ACKNOWLEDGMENTS

We thank Dr. Qingjun Wu from Institute of Vegetables and Flowers, CAAS for kindly supplying the whitheflies *Bemisia tabaci*. We also thank Dr. Xiaofei Cheng from Northeast Agricultural University for critical reading of the manuscript.

## REFERENCES

Arguello-Astorga, G.R., and Ruiz-Medrano, R. (2001). An iteron-related domain is associated to Motif 1 in the replication proteins of geminiviruses: identification of potential interacting amino acid-base pairs by a comparative approach. Arch. Virol. 146:1465–1485. 10.1007/s007050170072.

Cui, X., Tao, X., Xie, Y., Fauquet, C.M., and Zhou, X. (2004). A DNA beta associated with tomato yellow leaf curl China virus is required for symptom induction. J. Virol. 78:13966–13974.

Duffy, S., and Holmes, E.C. (2008). Phylogenetic evidence for rapid rates of molecular evolution in the single-stranded DNA begomovirus tomato yellow leaf curl virus. J. Virol. 82:957–965.

Hanley-Bowdoin, L., Bejarano, E.R., Robertson, D., and Mansoor, S. (2013). Geminiviruses: masters at redirecting and reprogramming plant processes. Nat. Rev. Microbiol. 11:777–788.

Harbach, C.J., Allen, T.W., Bowen, C.R., Davis, J.A., Hill, C.B., Leitman, M., Leonard, B.R., Mueller, D.S., Padgett, G.B., Phillips, X.A., et al. (2016). Delayed senescence in soybean: terminology, research update, and survey results from growers. Plant Health Progress 17:76–83.

Li, K., Zhang, X.X., Guo, J.Q., Penn, H., Wu, T.T., Li, L., Jiang, H., Chang, L.D., Wu, C.X., and Han, T.F. (2019). Feeding of *Riptortus pedestris* on soybean plants, the primary cause of soybean staygreen syndrome in the Huang-Huai-Hai river basin. Crop J. 7:360–367.

Savary, S., Willocquet, L., Pethybridge, S.J., Esker, P., McRoberts, N., and Nelson, A. (2019). The global burden of pathogens and pests on major food crops. Nat. Ecol. Evol. 3:430–439.

Veena, V., and Taylor, C.G. (2007). Agrobacterium rhizogenes: recent developments and promising applications. In Vitro Cell. Dev. Biol. Plant 43:383–403.

Xu, C., Han, T., and Wu, C. (2019). Discussion on the causes of staygreen syndrome for summer soybean and its preventive methods in the HuangHuai-Hai Region. Soybean Sci. Technol. 3:22–28 (in Chinese with English abstract).

Zhang, X.X., Wang, M., Wu, T.T., Wu, C.X., Jiang, B.J., Guo, C.H., and Han, T.F. (2016). Physiological and molecular studies of staygreen caused by pod removal and seed injury in soybean. Crop J. 4:435–443.

