## Supplementary material for "An intergeneric recombinant geminivirus causes soybean stay-green disease": All supplemental materials

### Supplementary Information

#### Supplemental Methods

##### Cloning, plasmids constructions and infectivity

The full-length SoSGV genome of were amplified using a pair of back-to-back primer P325 (5'-CAAGCTTGCATGCCTGCAGGTCGACGCTGTTAAGCGCCTTGGCGTAAGC-3', containing a *Sal*I site) and P326 (5'-CGAGCTCGGTACCCGGGGATCCGGCAGTAAGTCAGAGGCTCTTAA-3', containing a *Bam*H1 site) designed from assembled contigs using small RNAs from SoSGV infected soybean. This contig was located in the C1 region of SoSGV. DNA from SoSGV-infected soybeans was extracted by using CTAB method based extraction procedure (Murray and Thompson, 1980).

To construct an infectious clone of SoSGV, DNA from YY1 (collected from Yuanyang, Anhui province) was used as template and primers P325/P326 were used to amplify the first copy of full-length genome of SoSGV, and the PCR product was inserted into the pGEM-T vector (Promega, USA) to produce p1X-SoSGV. p1X-SoSGV was digested with *Sal*I and *Bam*HI and then was introduced into the binary vector pBinPLUS (Cui et al., 2005) to generate pBinPLUS-1xSoSGV (pBin1x-SoSGV). Next, the second copy of SoSGV was cloned by using primer P326 and P327 (5'-ATTAAGAGCCTCTGACTTACTGCCGCTGTTAAGCGCCTTGGCGTAAGC-3', containing *Bam*H1 site), and the PCR product was inserted into *Bam*H1 digested pBinPLUS-1xSoSGV vector to produce pBinPLUS-2xSoSGV (pBin2x-SoSGV). The pBlin2x-SoSGV was electro-incorporated into *A. tumefaciens* GV3101 and *A. rhizogenes* strain K599, respectively, for inoculation.

Infectivity of the cloned pBin2x-SoSGV was assessed by *Agrobacterium*-mediated inoculation as described previously (Cui, Tao et al. 2004). GV3101-

mediated inoculations onto N. *benthamiana*, *N. tabacum (*K326), *N. glutinosa* and Datura plants were conducted as described previously for other plant viruses (Briddon et al., 1989; Cui et al., 2005). For *A. rhizogenes* strain K599-mediated inoculation, we used hairy root transformation as described by Fan et al with minor modifications (Fan, Zhang et al. 2020). First, the primary root was cut from 8-day-old healthy seedlings with a sterile pair of scissors. In the inoculation medium, the top of hypocotyl was diagonally cut by sterile scalpel by 0.7-1cm. The slant cut of partial hypocotyl left was scraped on the plate grown K599 agrobacterium containing p2x-SoSGV and then directly planted into the pot with wet sterile vermiculite. Each plant was irrigated with 5 mL of treated *A. rhizogenes* liquid containing p2x-SoSGV plasmid ($OD_{600}$ = 1.0), and hairy root could be induced on inoculated sites *in vivo*. Treated plants were covered with a sterile plastic cage for 2 to 3 weeks before transferring to the soil-filled bowls. All these plants were maintained in insect proof cages in greenhouse and were monitored for up to 60 days for the appearance of symptoms. The plant leaves were subjected to PCR and Western blot at 4 wpi.

**Immunoblotting and antibody generation**

Protein extraction was performed as described previously (Xu, Ju et al. 2018) unless otherwise stated. The rabbit polyclonal antibody against SoSGV coat protein (CP) was generated in our lab. For immunoblotting, the membranes were detected with primary anti-CP antibodies (Sigma, St. Louis, USA), and followed with the corresponding secondary antibody (Bio-Rad, Hercules, USA) coupled with horseradish peroxidase. Blotted signal was visualized ECL using chemiluminescence according to the manufacturer's manual (Bio-Rad).

**Transmission assay**

Whiteflies (*Bemisia tabaci*, biotype Q) were reared in cotton plants in a cage. The cages are kept in the chamber at 26-28 °C and 65% relative humidity under 16/8 h day/night conditions. Transmission of progeny virus was assessed by caging groups of whitefiles on soybean plants infected with SoSGV by K599 *Agrobacterium*-mediated inoculation. A total of 500 healthy whitefly individuals reared on cotton were introduced to SoSGV-infected soybean source plants for 48-hr acquisition access period (AAP). Ten SoSGV-harbouring whiteflies were then transferred to 25 caged healthy soybeans (cotyledonary leaf stage) for 48-h inoculation access period (IAP). Following inoculation, the test plants were sprayed with an insecticide and kept in an insect-proof greenhouse pending the appearance of symptoms up to 60 days. Infection of the inoculated plants was tested by using PCR.

To examine seed-borne capacity of SoSGV, a random sample of seeds were collected from the SoSGV-infected soybeans. These seeds were first washed with stilled water followed by surface-sterilized with 75% ethanol for 10 min and 1.5% sodium hypochlorite for 20 min. Then the seed was used for extraction of DNA and test for the presence of SoSGV. In grow-out test, 100 seeds from SoSGV infected soybeans and 50 seeds from healthy plants were grown for gemination. All these germinated plants were maintained in insect proof cages in glass house and evaluated up to 60 days after emergence. True leaves were collected 30 days after planting and the leaves were subjected to DNA extraction and PCR detection using a SoSGV-CP primer (CP-F: 5'-ATGGATTACAGCAGGAAGAGG-3'; CP-R 5'-TTACAATTTGCTCTTGAAATACGT-3'). To test whether SoSGV could enter into endosperm and embryo, seeds collected from infected plants were first surface-sterilized and then dissected into seed coat, endosperm and embryo. Twenty seed coat, endosperm and embryo were subjected to DNA extraction by CTAB method. The extracted DNA from these tissues were subjected to PCR with SoSGV-CP primer.

**Phylogenetic, Recombination and Evolutionary Dynamics analysis**

Using the samples collected from Huang-Huai-Hai region in China, 27 genome-wide sequences were cloned and sequenced. Whole genome sequences of representative recognized members of each of the fourteen genera comprising the family *Geminiviridae* and 28 C1 (Rep) amino-acid sequences of representative recognized members of begomoviruses were retrieved from NCBI (http://www.ncbi.nlm.nih.gov/). Sequences were aligned by ClustalW implemented in MEGA X and phylogenetic trees were constructed by the ML method using IQ-TREE V2.0 (Kumar et al., 2016; Minh et al., 2020). One thousand replications were analyzed to enhance robustness. Recombination events were searched with SoSGV genome using complete genome sequences representing the major lineages of geminiviruses showing the highest similarities; the analysis was performed with SimPlot 3.5.1 software. Recombination analyses were performed using default settings and all trees were modified by using Figtree software version 1.4.4.

MCC trees were inferred by BEAST (v1.10.4) (Drummond and Rambaut, 2007), with GTR+I+G nucleotide substitution model. The relaxed uncorrected log-normal was specified as the molecular clock model. Convergence of each parameter was accepted with ESS > 200 in Tracer v1.7. Independent run was combined with LogCombiner v1.10.4 and the MCC tree was extracted with TreeAnnotator v1.10.4. Additionally, to identified Rep genotypes, three distinct methods were used including: MCC tree using BEAST (v1.10.4.4) (Drummond and Rambaut, 2007), ML tree using IQ-TREE V2.0 (Stamatakis, 2014),and NJ tree using MEGA X (Kumar et al., 2018). The p-distance methods were used to infer the NJ tree with 1000 bootstraps replication.

The detection of selection on the complete coding sequences of all SoSGV ORFs was performed using DATAMONKEY (http://www.datamonkey.org/). The methods used to investigate positive codon sites included SLAC, FEL, FUBAR, MEME (Murrell et al., 2012; 2013; Kosakovsky Pond and Frost, 2005); the branch site REL and the GA-branch site models were chosen to determine the selection pressure on the individual branches (Kosakovsky Pond and Frost, 2005b; Kosakovsky Pond et al., 2011). Methods with $p < 0.05$ in SLAC, FEL and MEME and the posterior probability >0.9 in FUBAR, were accepted, with at least three methods considered to be more conservative positive selection pressure.

#### Reference:


**Briddon, R. W., Watts, J., Markham, P. G. andStanley, J.** (1989). The coat protein of beet curly top virus is essential for infectivity. Virology **172**(2): 628-633.

**Cui, X., Li, G., Wang, D., Hu, D., and Zhou, X.** (2005). A Begomovirus DNAbeta-encoded protein binds DNA, functions as a suppressor of RNA silencing, and targets the cell nucleus. J. Virol. **79**(16): 10764-10775.

**Cui, X., Tao, X., Xie, Y., Fauquet, C. M., and Zhou, X.** (2004). A DNAβ associated with tomato yellow leaf curl China virus is required for symptom induction. J. Virol. **78**(24): 13966-13974.

**Drummond, A. J., and Rambaut, A.** (2007). BEAST: Bayesian evolutionary analysis by sampling trees. BMC Evol. Biol. **7**: 214.

**Fan, Y. L., Zhang, X. H., Zhong, L. J., Wang, X. Y., Jin L. S., and Lyu, S. H.** (2020). One-step generation of composite soybean plants with transgenic roots by *Agrobacterium rhizogenes*-mediated transformation. BMC Plant Biol. **20**(1): 208.

**Kosakovsky Pond, S. L., Frost, S. D.** (2005). A simple hierarchical approach to modeling distributions of substitution rates. Mol. Biol. Evol. **22**(2): 223-234.

**Kosakovsky Pond, S. L., Frost, S. D.** (2005b). A genetic algorithm approach to detecting lineage-specific variation in selection pressure. Mol. Biol. Evol. **22**(3): 478-485.

**Kosakovsky Pond, S. L., Murrell, B., Fourment, M., Frost S. D., Delport, W., Scheffler, K.** (2011). A random effects branch-site model for detecting episodic diversifying selection. Mol. Biol. Evol. **28**(11): 3033-3043.

**Kumar, S., Stecher, G., Li M., Knyaz C., and Tamura K**. (2018). MEGA X: Molecular evolutionary genetics analysis across computing platforms. Mol. Biol. Evol. **35**: 1547-1549.

**Minh, B. Q., Schmidt, H. A., Chernomor, O., Schrempf, D., Woodhams, M. D., von Haeseler A., and Lanfear, R.** (2020). IQ-TREE 2: New models and efficient methods for phylogenetic inference in the genomic era. Mol. Biol. Evol. **37**(5): 1530-1534.

**Murray, M. G., and Thompson, W. F.** (1980). Rapid isolation of high molecular weight plant DNA. Nucleic Acids Res. **8**(19): 4321-4325.

**Murrell, B., Moola, S., Mabona, A., Weighill, T., Sheward, D., Kosakovsky Pond, S. K., Scheffler, K.** (2013). FUBAR: a fast, unconstrained bayesian approximation for inferring selection. Mol. Biol. Evol. **30**(5): 1196-205.

**Murrell, B., Wertheim, J. O., Moola, S., Weighill, T., Scheffler, K., Kosakovsky Pond, S. L.** (2012). Detecting individual sites subject to episodic diversifying selection. PLoS Genet. **8**(7): e1002764.

**Stamatakis, A.** (2014). RAxML version 8: a tool for phylogenetic analysis and post-analysis of large phylogenies. Bioinformatics **30**(9): 1312-1313.

**Xu, Y., Ju, H. J., DeBlasio, S., Carino, E. J., Johnson, R., MacCoss, M. J., Heck, M., Miller, W. A., and Gray, S. M.** (2018). A stem-loop structure in potato leafroll virus open reading frame 5 (ORF5) is essential for readthrough translation of the coat protein ORF stop codon 700 bases upstream. J. Virol. **92**(11).

#### Supplemental Tables

**Table S1. The ORFs of SoSGV and the related geminiviruses with the highest sequence identities.**

| ORF | Locus | nt length | Virus with highest nt similarity (Percentage) | Virus with highest AA similarity | Genus of the most similar virus |
|---|---|---|---|---|---|
| V1 (CP) | 270-1055 | 786 | Nd | SPSV1 (29.6%) | *Mastrevirus* |
| | | | | CYV (27.2) | *Mastrevirus* |
| V2 | 146-457 | 312 | JMAV (51.8) | JMAV (25.5) | *Maldovirus* |
| C1 | 1539-2624 | 1086 | TLCJV (84.1) | TLCJV (84.5) | *Begomovirus* |
| C2 | 1220-1636 | 417 | JMAV (63.4) | JMAV (44.4) | *Maldovirus* |
| C3 | 1069-1524 | 456 | JMAV (63.4) | JMAV (44.8) | *Maldovirus* |
| C4 | 2177-2467 | 291 | TLCJV (92) | AYVV (82) | *Begomovirus* |

Chickpea yellows virus, CYV; Juncus maritimus associated virus, JMAV; Tomato leaf curl Java virus-[Ageratum], TLCJV; sweet potato symptomless virus 1, SPSV1; Nd, not detected.

**Table S2. Agroinoculation infectivity assay and host range tests**

| Plant species | No. plants infiltrated | Agrobacteria used | No. plants Systemically infected | Seedlings symptom | Infection efficiency (%) |
|---|---|---|---|---|---|
| ***N. benthamiana*** | 30 | GV3101[a] | 30 | Leaf crinkle, dwarf | 100% |
| ***N. tabacum* (K326)** | 30 | GV3101[a] | 28 | Leaf crinkle, dwarf | 93.3% |
| ***N. glutinosa*** | 30 | GV3101[a] | 27 | Leaf crinkle, | 100% |
| ***Datura stramonium*** | 30 | GV3101[a] | 27 | Leaf crinkle | 90% |
| ***S. lycopersicum* Moneymaker** | 30 | GV3101[a] | 0 | nd[c] | 0% |
| ***S. lycopersicum* M82** | 30 | GV3101[a] | 0 | nd | 0% |
| ***Glycine max* Williams 82** | 30 | GV3101[a] | 0 | nd | 0% |
| ***Glycine max* Wanhua 518** | 30 | Gv3101[a] | 0 | nd | 0% |
| ***Glycine max* Williams 82** | 30 | K599[b] | 27 | Leaf crinkle | 90% |
| ***Glycine max* Wanhua 518** | 30 | K599[b] | 30 | Leaf crinkle | 100% |
| **Shanning** | 30 | K599[b] | 29 | Leaf crinkle | 96.7% |
| **Nannong1138-2** | 30 | K599[b] | 30 | Leaf crinkle | 100% |

[a] GV3101-mediated syringe injection; [b] *A. rhizogenes* strain K599-mediated inoculation; [c] not detected.

**Table S3. Whitefly transmission assay**

| | No. of plant infected with SoSGV/no. infested with whiteflies | |
|---|---|---|
| | **test 1** | **test 2** |
| **SoSGV** | **0/15** | **0/30** |

**Table S4. Seed transmission test**

| Seed origin/part | Test Number | Positive number |
|---|---|---|
| **Natural infection** | | |
| whole seed | 120 | 108/108 |
| seed coat | 108 | 108/108 |
| cotyledon | 108 | 0/108 |
| embryo | 106 | 0/106 |
| grow-out leaves | 100 | 0/93 |
| **Experimental infection (K599)** | | |
| whole seed | 50 | 50/50 |
| seed coat | 50 | 50/50 |
| cotyledon | 50 | 0/50 |
| embryo | 48 | 0/48 |
| grow-out leaves | 27 | 0/27 |

**Table S5. Twenty-seven SoSGV whole genome sequences deposited into NCBI**

| Name | GenBank No. | Collected location | Collected Year | No. in Fig.1 |
|---|---|---|---|---|
| XX-1_2019 | OM145983 | Xinxiang, Henan | 2019 | 1 |
| XX-2_2019 | OM145984 | Xinxiang, Henan | 2019 | 2 |
| XX-3_2019 | OM145985 | Xinxiang, Henan | 2019 | 3 |
| YY-1_2019 | OM145986 | Yuanyang, Henan | 2019 | 4 |
| YY-2_2019 | OM145987 | Yuanyang, Henan | 2019 | 5 |
| YY-3_2019 | OM145988 | Yuanyang, Henan | 2019 | 6 |
| ZZ-1_2019 | OM145989 | Zhengzhou, Henan | 2019 | 7 |
| ZZ-1_2021 | OM145990 | Zhengzhou, Henan | 2021 | 8 |
| ZZ-2_2019 | OM145991 | Zhengzhou, Henan | 2019 | 9 |
| ZZ-2_2021 | OM145992 | Zhengzhou, Henan | 2021 | 10 |
| ZZ-3_2019 | OM145993 | Zhengzhou, Henan | 2019 | 11 |
| ZZ-3_2021 | OM145994 | Zhengzhou, Henan | 2021 | 12 |
| ZZ-4_2019 | OM145995 | Zhengzhou, Henan | 2019 | 13 |
| ZZ-5_2019 | OM145996 | Zhengzhou, Henan | 2019 | 14 |
| ZZ-6_2019 | OM145997 | Zhengzhou, Henan | 2019 | 15 |
| ZZ-7_2019 | OM145998 | Zhengzhou, Henan | 2019 | 16 |
| ZZ-8_2019 | OM145999 | Zhengzhou, Henan | 2019 | 17 |
| ZZ-9_2019 | OM146000 | Zhengzhou, Henan | 2019 | 18 |
| ZZ-10_2019 | OM146001 | Zhengzhou, Henan | 2019 | 19 |
| ZZ-11_2019 | OM146002 | Zhengzhou, Henan | 2019 | 20 |
| ZZ-12_2019 | OM146003 | Zhengzhou, Henan | 2019 | 21 |
| ZZ-13_2019 | OM146004 | Zhengzhou, Henan | 2019 | 22 |
| ZZ-14_2019 | OM146005 | Zhengzhou, Henan | 2019 | 23 |
| SZ-1_2019 | OM146006 | Suzhou, Anhui | 2019 | 24 |
| SZ-1_2021 | OM146007 | Suzhou, Anhui | 2021 | 25 |
| SZ-3_2021 | OM146008 | Suzhou, Anhui | 2021 | 26 |
| SZ-4_2021 | OM146009 | Suzhou, Anhui | 2021 | 27 |

**Table S6. Selection analysis of SoSGV ORFs**

| FEL | p | MEME | p | FUBAR | Posterior probability |
|---|---|---|---|---|---|
| C1, 219 | 0.09 | C1, 219 | 0.06 | C1, 219 | 0.942 |

#### Supplemental Figures

**Fig. S1 Soybean plants with stay-green symptoms from fields and structure analysis of SoSGV IR region and profile of SoSGV-derived sRNA.** (*A*), Samples collected from field in the Huang-Huai-Hai region of China with typical stay-green symptom of whole plant; Left: healthy plant; Right: SoSGV-infected plant (*B*), Soybean pods of healthy (top) and stay-green (bottom) samples from full angle of view. Top: pods from healthy plant; Bottom: pods from SoSGV-infected plant. (*C*), Soybean seeds from healthy (top) and the stay-green field samples (bottom). (*D*) The stem-loop in the IR of SoSGV and with conserved geminiviral nonanucleotide "TAATATTAC" as indicated by arrows. (*E*) iterated DNA motif (iteron) and the corresponding iteron-related domain (IRD) in SoSGV IR region. (*F*) Length distributions of vsiRNA sequences originated from SoSGV genome (YY1). (*G*) Profile of SoSGV-derived vsiRNAs from along the SoSGV genome. The horizontal axis represents the SoSGV genomes. Numbers on the y axis correspond to the amount of vsiRNAs mapped to the SoSGV genomic (+) or antigenomic (-) sequences. "All" represents the total small RNAs from SoSGV and "21" represents the 21-nt vsiRNAs from SoSGV.

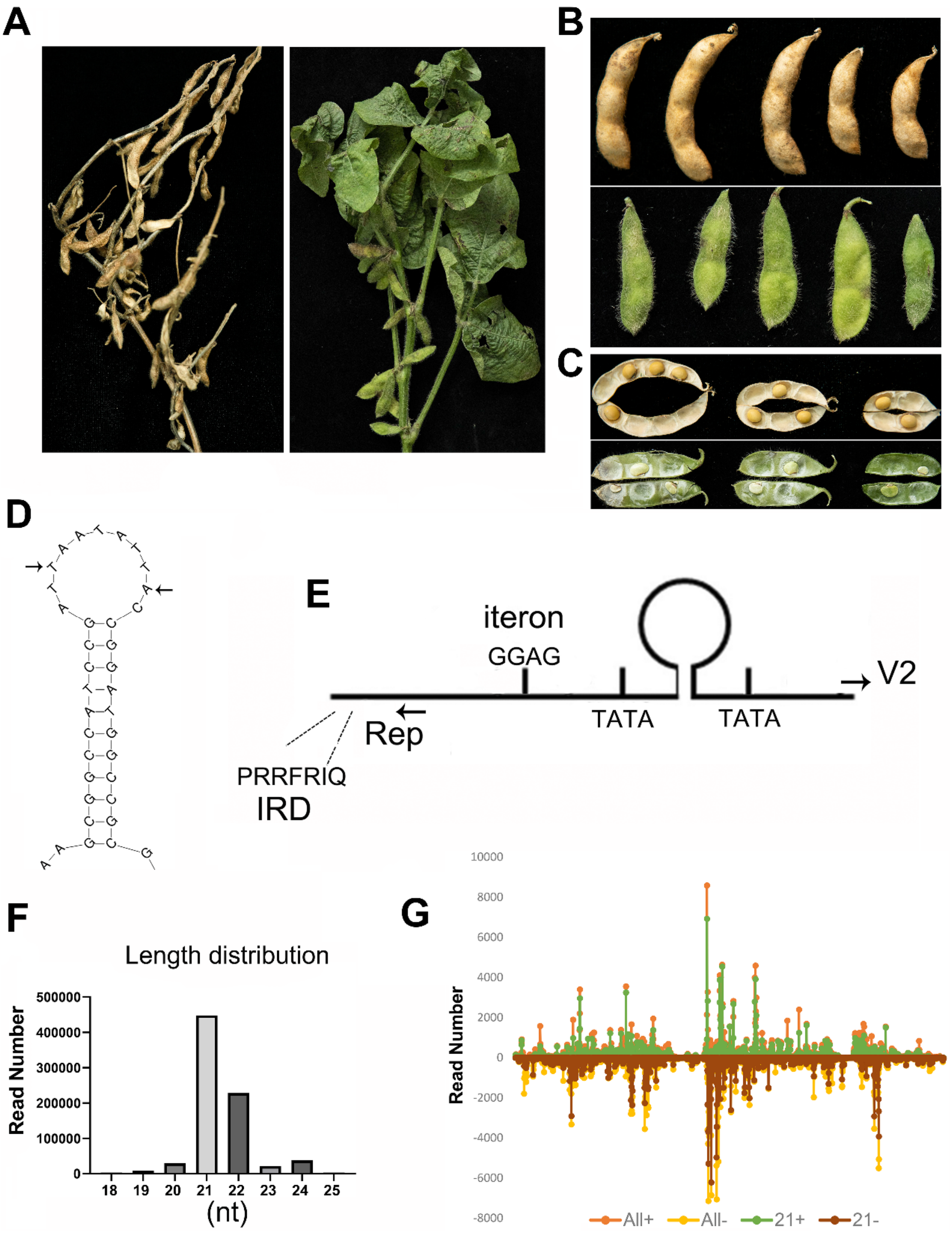
A
B
C
D
E
iteron
GGAG
V2
TATA
TATA
Rep
PRRFRIQ
IRD
F
Length distribution
Read Number
500000
400000
300000
200000
100000
0
18
19
20
21
22
23
24
25
(nt)
G
Read Number
10000
8000
6000
4000
2000
0
-2000
-4000
-6000
-8000
All+
All-
21+
21-

**Fig. S2 Phylogenetic tree based on full genome sequences of SoSGV and selected geminiviruses.** The tree was constructed using the maximum-likelihood (ML) method in the MEGA X software. Whole genome sequences of representative recognized members of each of the fourteen genera comprising the family *Geminiviridae* were retrieved from NCBI. The accession number was marked and different color represents viruses in the same genera of *Geminiviridae* family.

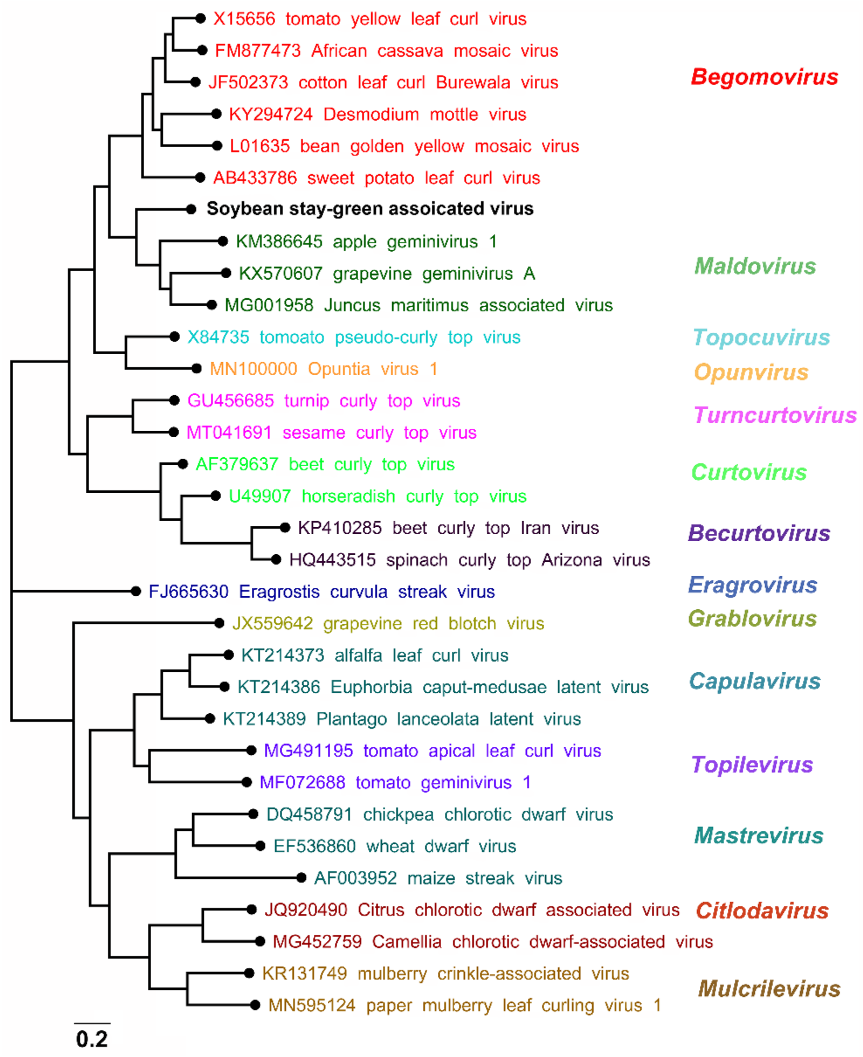

**Fig. S3 Genome recombinations in SoSGV.** (*A*) Whole genome sequence similarity plot of geminiviruses including Juncus maritimus associated virus (isolate 13-FMN-1), apple geminivirus 1 (NC_026760) and tomato leaf curl Java virus-[Ageratum] (AB162141.1) using the SoSGV as a query. The analysis was performed using Simplot, with a window size of 200 bp and a step size of 20 bp. The horizontal axis represents the SoSGV genomes. Numbers on the y axis correspond to the similarity value. (*B*-*G*) Phylogenetic analysis of amino acids of SoSGV ORFs and representative geminiviruses. (*B*) the V1 gene, (*C*) the V2 gene, (*D*) the C1 gene, (*E*) the C2 gene, (*F*) the C3 gene and (*G*) the C4 gene. Phylogenetic analysis was performed by the ML method using IQ-TREE V2.0 with 1000 bootstrap replicates. Branch lengths are scaled according to the number of nucleotide substitutions per site. SoSGV is indicated by red text in each figure.

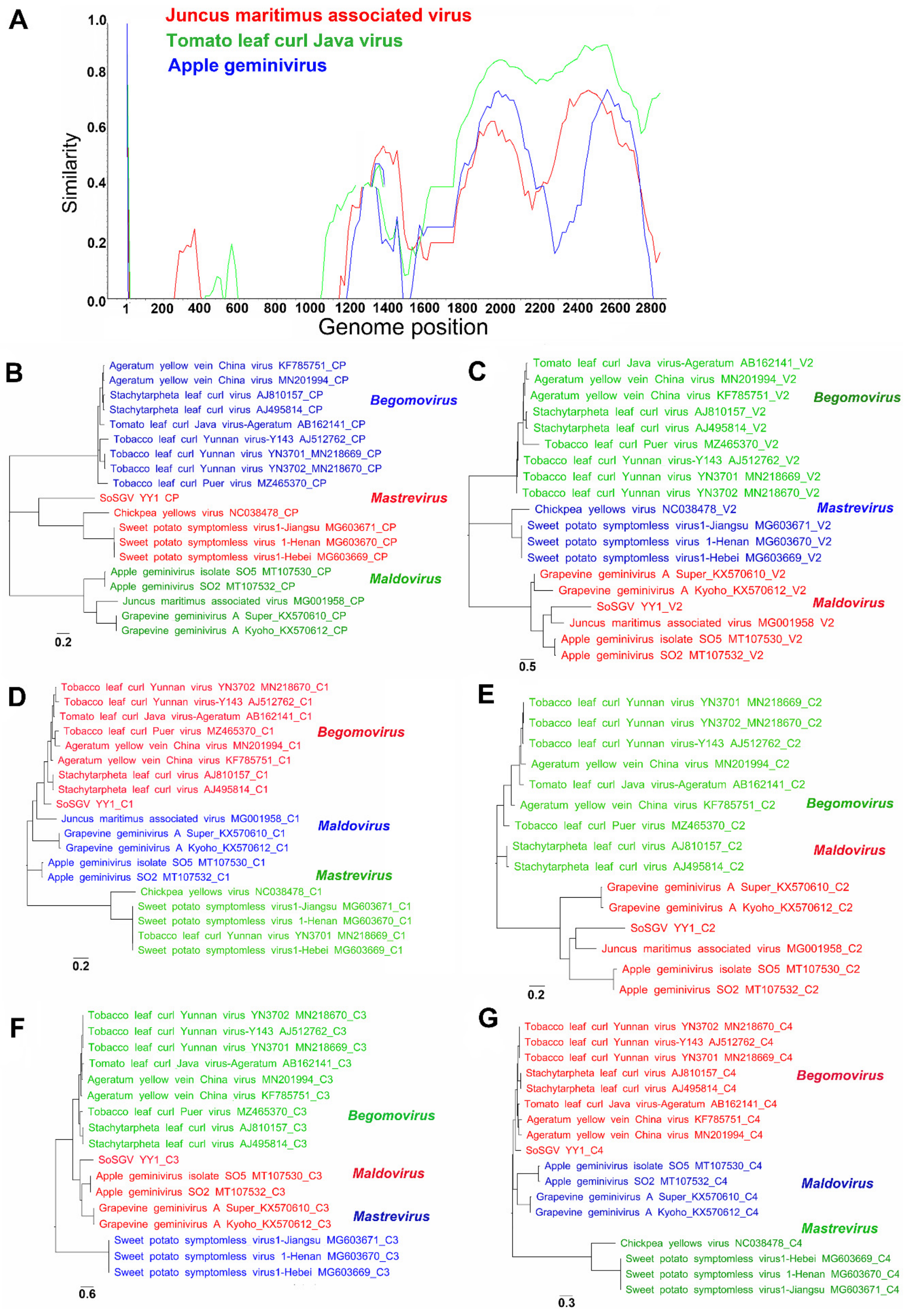

A
Juncus maritimus associated virus
Tomato leaf curl Java virus
Apple geminivirus
Similarity
0.0
0.2
0.4
0.6
0.8
1.0
1
200
400
600
800
1000
1200
1400
1600
1800
2000
2200
2400
2600
2800
Genome position
B
Ageratum yellow vein China virus KF785751_CP
Ageratum yellow vein China virus MN201994_CP
Stachytarpheta leaf curl virus AJ810157_CP
Stachytarpheta leaf curl virus AJ495814_CP
Tomato leaf curl Java virus-Ageratum AB162141_CP
Tobacco leaf curl Yunnan virus-Y143 AJ512762_CP
Tobacco leaf curl Yunnan virus YN3701_MN218669_CP
Tobacco leaf curl Yunnan virus YN3702_MN218670_CP
Tobacco leaf curl Puer virus MZ465370_CP
SoSGV YY1 CP
Chickpea yellows virus NC038478_CP
Sweet potato symptomless virus1-Jiangsu MG603671_CP
Sweet potato symptomless virus 1-Henan MG603670_CP
Sweet potato symptomless virus1-Hebei MG603669_CP
Apple geminivirus isolate SO5 MT107530_CP
Apple geminivirus SO2 MT107532_CP
Juncus maritimus associated virus MG001958_CP
Grapevine geminivirus A Super_KX570610_CP
Grapevine geminivirus A Kyoho_KX570612_CP
Begomovirus
Mastrevirus
Maldovirus
0.2
C
Tomato leaf curl Java virus-Ageratum AB162141_V2
Ageratum yellow vein China virus MN201994_V2
Ageratum yellow vein China virus KF785751_V2
Stachytarpheta leaf curl virus AJ810157_V2
Stachytarpheta leaf curl virus AJ495814_V2
Tobacco leaf curl Puer virus MZ465370_V2
Tobacco leaf curl Yunnan virus-Y143 AJ512762_V2
Tobacco leaf curl Yunnan virus YN3701 MN218669_V2
Tobacco leaf curl Yunnan virus YN3702 MN218670_V2
Chickpea yellows virus NC038478_V2
Sweet potato symptomless virus1-Jiangsu MG603671_V2
Sweet potato symptomless virus 1-Henan MG603670_V2
Sweet potato symptomless virus1-Hebei MG603669_V2
Grapevine geminivirus A Super_KX570610_V2
Grapevine geminivirus A Kyoho_KX570612_V2
SoSGV YY1_V2
Juncus maritimus associated virus MG001958 V2
Apple geminivirus isolate SO5 MT107530_V2
Apple geminivirus SO2 MT107532_V2
Begomovirus
Mastrevirus
Maldovirus
0.5
D
Tobacco leaf curl Yunnan virus YN3702 MN218670_C1
Tobacco leaf curl Yunnan virus-Y143 AJ512762_C1
Tomato leaf curl Java virus-Ageratum AB162141_C1
Tobacco leaf curl Puer virus MZ465370_C1
Ageratum yellow vein China virus MN201994_C1
Ageratum yellow vein China virus KF785751_C1
Stachytarpheta leaf curl virus AJ810157_C1
Stachytarpheta leaf curl virus AJ495814_C1
SoSGV YY1_C1
Juncus maritimus associated virus MG001958_C1
Grapevine geminivirus A Super_KX570610_C1
Grapevine geminivirus A Kyoho_KX570612_C1
Apple geminivirus isolate SO5 MT107530_C1
Apple geminivirus SO2 MT107532_C1
Chickpea yellows virus NC038478_C1
Sweet potato symptomless virus1-Jiangsu MG603671_C1
Sweet potato symptomless virus 1-Henan MG603670_C1
Tobacco leaf curl Yunnan virus YN3701 MN218669_C1
Sweet potato symptomless virus1-Hebei MG603669_C1
Begomovirus
Maldovirus
Mastrevirus
0.2
E
Tobacco leaf curl Yunnan virus YN3701 MN218669_C2
Tobacco leaf curl Yunnan virus YN3702_MN218670_C2
Tobacco leaf curl Yunnan virus-Y143 AJ512762_C2
Ageratum yellow vein China virus MN201994_C2
Tomato leaf curl Java virus-Ageratum AB162141_C2
Ageratum yellow vein China virus KF785751_C2
Tobacco leaf curl Puer virus MZ465370_C2
Stachytarpheta leaf curl virus AJ810157_C2
Stachytarpheta leaf curl virus AJ495814_C2
Grapevine geminivirus A Super_KX570610_C2
Grapevine geminivirus A Kyoho_KX570612_C2
SoSGV YY1_C2
Juncus maritimus associated virus MG001958_C2
Apple geminivirus isolate SO5 MT107530_C2
Apple geminivirus SO2 MT107532_C2
Begomovirus
Maldovirus
0.2
F
Tobacco leaf curl Yunnan virus YN3702 MN218670_C3
Tobacco leaf curl Yunnan virus-Y143 AJ512762_C3
Tobacco leaf curl Yunnan virus YN3701 MN218669_C3
Tomato leaf curl Java virus-Ageratum AB162141_C3
Ageratum yellow vein China virus MN201994_C3
Ageratum yellow vein China virus KF785751_C3
Tobacco leaf curl Puer virus MZ465370_C3
Stachytarpheta leaf curl virus AJ810157_C3
Stachytarpheta leaf curl virus AJ495814_C3
SoSGV YY1_C3
Apple geminivirus isolate SO5 MT107530_C3
Apple geminivirus SO2 MT107532_C3
Grapevine geminivirus A Super_KX570610_C3
Grapevine geminivirus A Kyoho_KX570612_C3
Sweet potato symptomless virus1-Jiangsu MG603671_C3
Sweet potato symptomless virus 1-Henan MG603670_C3
Sweet potato symptomless virus1-Hebei MG603669_C3
Begomovirus
Maldovirus
Mastrevirus
0.6
G
Tobacco leaf curl Yunnan virus YN3702 MN218670_C4
Tobacco leaf curl Yunnan virus-Y143 AJ512762_C4
Tobacco leaf curl Yunnan virus YN3701 MN218669_C4
Stachytarpheta leaf curl virus AJ810157_C4
Stachytarpheta leaf curl virus AJ495814_C4
Tomato leaf curl Java virus-Ageratum AB162141_C4
Ageratum yellow vein China virus KF785751_C4
Ageratum yellow vein China virus MN201994_C4
SoSGV YY1_C4
Apple geminivirus isolate SO5 MT107530_C4
Apple geminivirus SO2 MT107532_C4
Grapevine geminivirus A Super_KX570610_C4
Grapevine geminivirus A Kyoho_KX570612_C4
Chickpea yellows virus NC038478_C4
Sweet potato symptomless virus1-Hebei MG603669_C4
Sweet potato symptomless virus 1-Henan MG603670_C4
Sweet potato symptomless virus1-Jiangsu MG603671_C4
Begomovirus
Maldovirus
Mastrevirus
0.3

**Fig. S4** ***A. tumefaciens*** **strain GV3101-mediated inoculation of infectious clone of SoSGV into** ***N. benthamiana*****,** ***N. glutinosa*** **and** ***Datura stramonium*** **plants.** The infectivity of the cloned genome of SoSGV by Agrobacterium (GV3101)-mediated injection to *N. benthamiana* (*A*), *N. glutinosa* (*B*) and *Datura stramonium* (*C*). Leaves were confirmed as systemically infected using PCR and Western blot. The number in the Figure represents SoSGV inoculation efficiency (systemically infected plants/total inoculated plants). Plasmid pBin2x-SoSGV was used as positive control in the PCR assay, and protein extract from SoSGV infected *N. benthamiana* plant was used as positive control in Western blot assay.

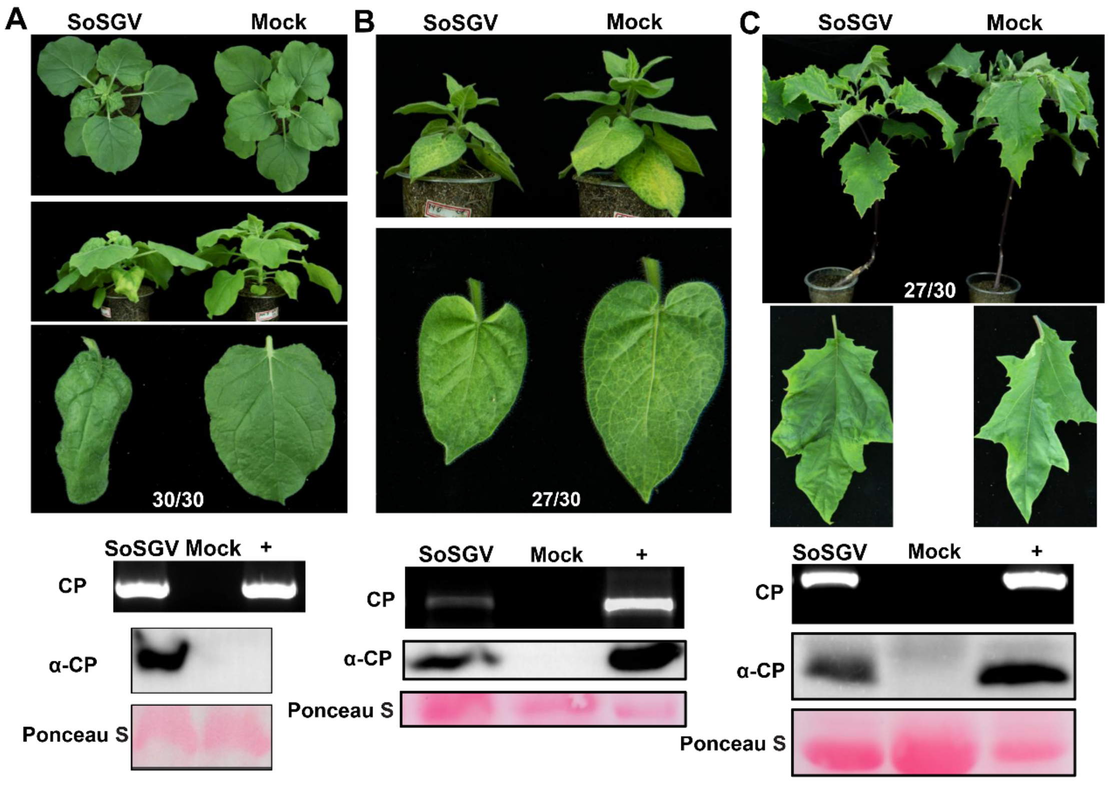

**Fig. S5** ***A. rhizogenes*** **strain K599-mediated inoculation of infectious clone of SoSGV and leaf crinkle symptom on infected soybean plants.** SoSGV inoculation of soybean cultivar Williams 82 (*A*) and Wanhua 518 (*B*) mediated by *A. rhizogenes* and the infections were confirmed by both PCR and Western blot. Leaf of soybean seedlings infected with cloned SoSGV by agroinoculation produces leaf crinkle symptom 1.5-month post inoculation. The number in the figure represents SoSGV inoculation efficiency (systemically infected plants/total inoculated plants).

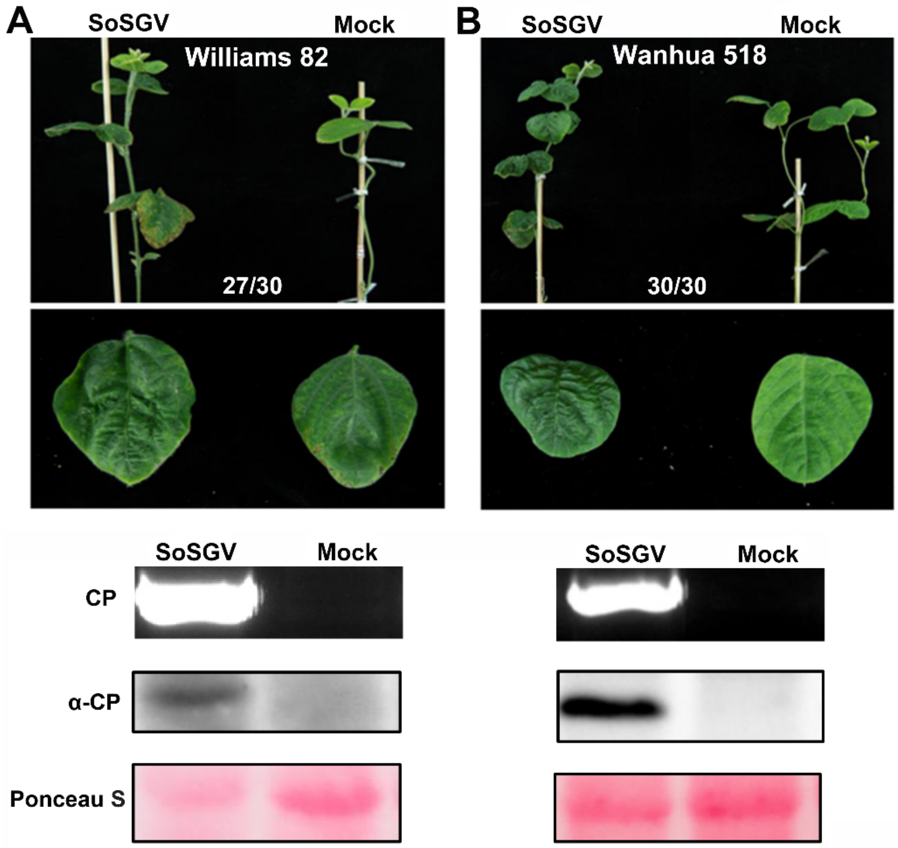

**Fig. S6 Inoculation of SoSGV infectious clone through *A. tumefaciens* strain GV3101-mediated inoculation causes *N. benthamiana* stay-green like symptoms.** Fifty days after inoculation of *N. benthamiana* plants by infectious clone of SoSGV, all mock inoculated plants (30/30) display mature seeds (*A*) and more yellowing leaves (*B*), whereas for SoSGV infection, the seed production are severely affected (*A*) and most of the leaves are still stay-green (*B*).

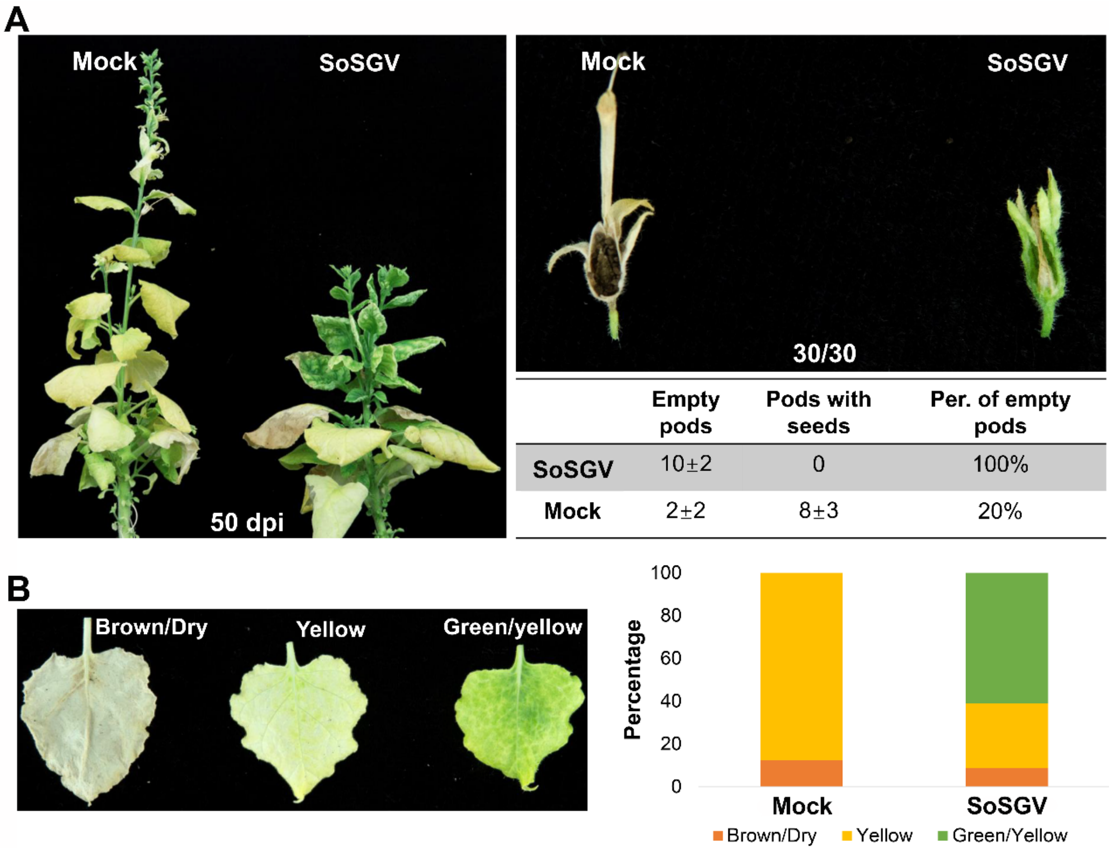


| | Empty pods | Pods with seeds | Per. of empty pods |
|---|---|---|---|
| SoSGV | 10±2 | 0 | 100% |
| Mock | 2±2 | 8±3 | 20% |

**Fig. S7 The abundance of SoSGV in soybean seeds compared with other plant tissues.** (*A*) The presence of SoSGV in seed of soybeans inoculated with SoSGV infectious clone through *A. rhizogenes* strain K599-mediated inoculation. (*B*) Semi-quantitative PCR was performed using CP primers, and the number of amplification cycles was 20. Plasmid pBin2x-SoSGV was used as positive control and soybean *tubulin* gene was used as an internal reference gene.

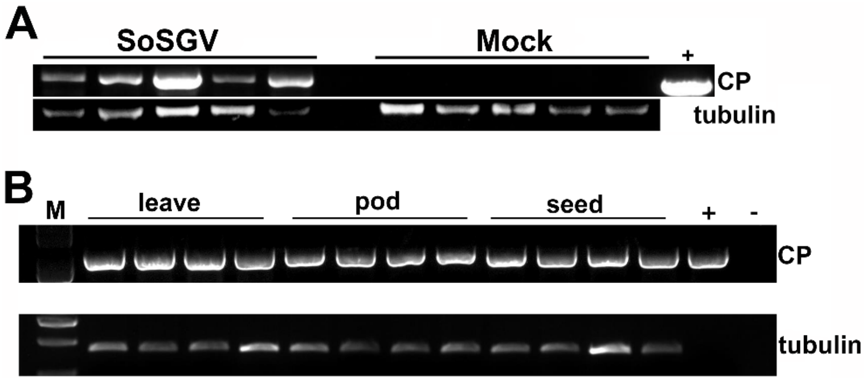


**Fig. S8 Locations in Huang-Huai-Hai region of China for samples collection and the epidemiological dynamics analysis of SoSGV.** MCC tree scaled to time using BEAST (v1.10.4) with the GTR+I+G substitution model and an uncorrected relaxed clock (lognormal) of SoSGV genomic sequences. Twenty-eight SoSGV genomic sequences including samples collected from Huang-Huai-Hai region of China and an isolate from South Korea were used. All the sequences have been deposited in the NCBI. The posterior displayed along each branch.

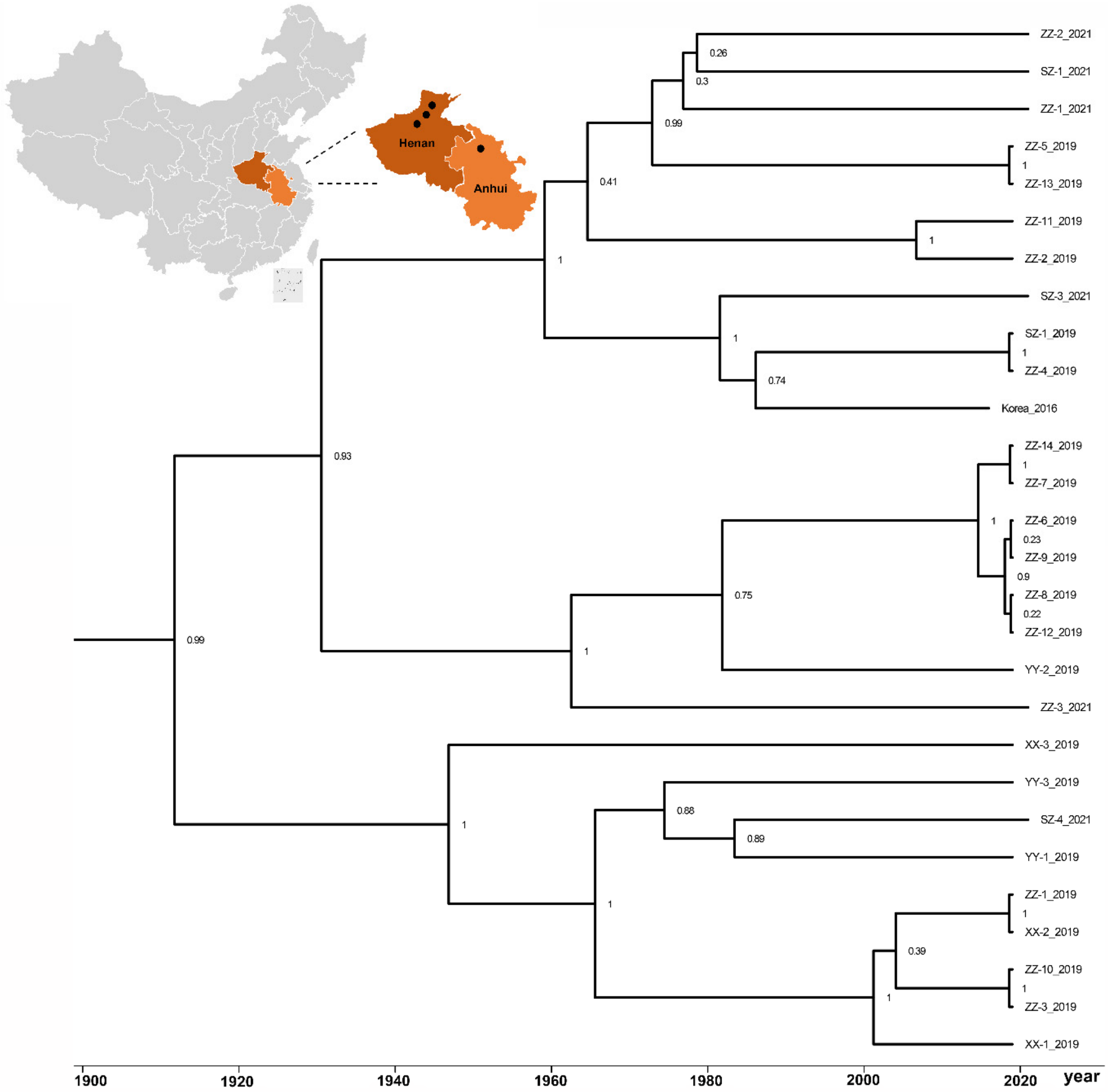


**Dataset 1. Estimates of evolutionary divergence among SoSGV isolates and ORF nt sequences.**

**Dataset 2. MCC file for analysis of molecular variability of SoSGV whole genome.**
